# Reproduction of a marine planktonic protist: Individual success versus population survival

**DOI:** 10.1101/2020.11.04.368100

**Authors:** Manuel F. G. Weinkauf, Michael Siccha, Agnes K. M. Weiner

**Affiliations:** Center for Marine Environmental Sciences (MARUM), Universität Bremen, Leobener Str. 8, 28359 Bremen, Germany; Department of Earth Sciences, Université de Genève, Rue des Maraîchers 13, 1205 Genève, Switzerland; Institute of Geology and Palaeontology, Univerzita Karlova, Albertov 2038/6, 128 43 Praha, Czech Republic

## Abstract

Understanding the biology of reproduction is important for retracing key evolutionary processes in organisms, yet gaining detailed insights often poses major challenges. Planktonic Foraminifera are globally distributed marine microbial eukaryotes and important contributors to the global carbon cycle. Their extant biodiversity shows restricted distribution patterns of some species, whereas others are cosmopolitan in the world ocean. Planktonic Foraminifera cannot be bred under laboratory conditions, and thus details of their life cycle remain incomplete. Solely the production of flagellated gametes has been observed and taken as an indication for an exclusively sexual reproduction. Yet, sexual reproduction by spawning of gametes in the open ocean relies on sufficient gamete encounters to maintain viable populations, which represents a problem for organisms that lack the means of active propulsion and are marked by low population densities. To increase knowledge on the reproductive biology of planktonic Foraminifera, we applied a dynamic, individual-based modelling approach with parameters based on laboratory and field observations to test if random gamete encounters under commonly observed population densities are sufficient for maintaining viable populations. We show that temporal synchronization and potentially spatial concentration of gamete release seems inevitable for maintenance of the population. We argue that planktonic Foraminifera optimized their individual reproductive success at the expense of community-wide gene flow, which may explain their high degree of diversity. Our modelling approach helps to illuminate foraminiferal population dynamics and to predict the existence of necessary reproduction strategies, which may be detected in future field experiments. This study therefore contributes to our understanding of plankton ecology and evolution and their reproductive strategies in the open ocean.

## 1 Introduction

The mode of reproduction exerts a strong influence on the evolution of organisms (Kondrashov 1988, Kirkwood and Rose 1991, Nilsson and Svensson 1996). In order to retrace processes such as adaptation, diversification, and speciation as well as population dynamics of organisms, it is necessary to first understand how they reproduce. Sexual reproduction is assumed to be an ancient trait that is widespread among eukaryotic taxa, including protists (Speijer et al. 2015, Lenormand et al. 2016, Mirzaghaderi and Hörandl 2016). Its existence has a major influence on populations, since gene flow alters genetic diversity and the potential for local adaptations and speciation.

In the open ocean, sexual reproduction confronts planktonic organisms with the problem of mate encounter to ensure sufficient reproductive success over wide geographical areas to maintain viable populations. While some plankton species are characterized by restricted distribution patterns with spatially limited gene flow (Casteleyn et al. 2010, Morard et al. 2011, Ujiié et al. 2012, Weiner et al. 2012, Godhe et al. 2013), there is also ample evidence for the existence of cosmopolitan species that maintain gene flow on a global scale (Lazarus et al. 1995, Renaud and Schmidt 2003, McManus and Woodson 2012, André et al. 2013, Weiner et al. 2014). Since planktonic organisms often show a patchy distribution in the open ocean with large variations in population densities, sexually reproducing species with wide dispersal potential must have developed efficient strategies to overcome the obstacle of mate encounter to maintain gene flow even across areas with low population densities. Such adaptive strategies include synchronization of reproduction in time and/or space (Bijma et al. 1990, Erez et al. 1991). efficient mate detection mechanisms through chemical communication (Blackburn et al. 1998, Fenchel 2001), or the production of a large number of motile gametes. While this problem affects all planktonic organisms living in the open ocean, it presents a major challenge for micro- and meso-zooplankton, such as marine planktonic protists. Due to their small size and limited means for active locomotion they are subject to passive dispersal by ocean currents, and oceanic turbulences may impede successful mate encounter and counteract adaptive strategies (Borgnino et al. 2019). Our knowledge on the reproduction of many protists in their natural habitat remains limited due to intrinsic difficulties of laboratory culturing. Consequently, many protist species are so far assumed to reproduce purely asexually (Coats and Heinbokel 1982, Weisse 2008). However, complex life cycles with an alternation of sexual and asexual stages occur in some protist groups (Goldstein 1997, Fiore-Donno et al. 2011), and others are even assumed to reproduce exclusively by sexual reproduction (Goldstein 2002).

Planktonic Foraminifera are a group of non-motile marine protists that produce calcitic shells and make up a large proportion of the meso-zooplankton in the world’s oceans (Schiebel 2002). Due to their fossil record, planktonic Foraminifera became widely used as proxy organisms in micropalaeontological and palaeoceanographic studies (Kučera 2007). Further, they represent excellent model organisms for studying the evolutionary history of marine protists, as their fossil record is nearly complete and reflects speciation and extinction events since their origin in the early Jurassic (Aze et al. 2011). However, full exploitation of their potential as model organisms in evolutionary studies is hampered by the fact that they cannot be cultured for an extended length of time under laboratory conditions and that our knowledge on their modes of reproduction remains scarce. Their complete life cycle has not yet been observed, though shortterm culturing experiments revealed the formation of flagellated gametes (Bé and Anderson 1976, Bé et al. 1977, Schiebel and Hemleben 2017). Adult individuals were reported to produce hundreds of thousands of biflagellated gametes that are released into the surrounding water (Bé and Anderson 1976). The formation of these gametes was taken as an indication for sexual reproduction, although fusion of the gametes and zygote formation have never been observed. Neither has self-fertilization ever been reported, and asexual reproduction was only detected in rare cases (Takagi et al. 2020). Consequently, planktonic Foraminifera are currently assumed to be mainly sexually reproducing, dioecious organisms (Schiebel and Hemleben 2017). This contrasts with benthic Foraminifera, which exhibit a wide range of reproductive modes amongst which a heterophasic life cycle alternating between sexual and asexual stages dominates (Goldstein 2002). Since planktonic species evolved from benthic ancestors (Darling et al. 1997, Hart et al. 2002), it was argued that switching to purely sexual reproduction represents an important step in the evolution from benthic to pelagic lifestyle (Kučera, et al. 2017). However, when considering the patchy occurrence of planktonic Foraminifera in the open ocean with very low population densities (on average 5–50 spec. m^−3^; Kučera et al. 2013, Meilland et al. 2019), the question arises on how viable populations can be maintained by sexual reproduction alone. To assure sufficient gamete encounters in the open ocean, the number of gametes produced per adult individual has to be adequately large, the survival time of the gametes long enough, and they have to disperse over large distances to meet each other. So far, it remains unknown if this can be achieved by random gamete encounters or if further adaptive strategies, such as temporal/spatial synchronization of reproduction or communication via chemical traits are necessary to enhance gamete encounter rates.

Considering the difficulties for maintaining viable populations in the open ocean environment, strong selective pressure exists in plankton organisms for reproductive strategies that enhance the chances of gamete encounter, which makes the existence of such adaptations in planktonic Foraminifera highly probable. Since observing plankton reproduction under natural conditions remains difficult, and laboratory culturing still faces obstacles, in this study we supplement already existing observations from field and laboratory experiments with mathematical modelling. Based on a spectrum of generally observed population densities and gamete numbers as well as observations on gamete size and speed we model the rates of gamete fusion by chance encounter. We further estimate the number of zygotes that must survive and reach a reproductive state to maintain the population. This allows us to evaluate if species of planktonic Foraminifera are able to maintain their populations in the open ocean by relying on sexual reproduction or if asexual reproduction plays a more important role for population growth than currently assumed. We further evaluate if strategies such as temporal and spatial synchronization are at play to assure successful reproduction.

## 2 Material and methods

We modelled planktonic Foraminifera reproduction in MatLab v. R2017b to test possible reproductive pathways that may occur in nature. An entirely realistic model, with several hundreds of thousands of particles interacting in a turbulent three-dimensional environment was computationally not feasible. Our model therefore comprises several parameterizations and models planktonic Foraminifera and their gametes in a laminar flow environment. The model consists of five modules: (1) Simulation of the distribution of gametes in space and time after gamete release; (2) simulation of several individual Foraminifera. in a quasi-laminar flow environment; (3) generation of spawning parameters for individual Foraminifera (e.g. time of gamete release, number of released gametes); (4) determination of probable encounters of gametes of different individuals; and (5) calculation of the number of fusions occurring based on the results of the previous module. The modular design allowed the re-use of experimental setups, so that for example the trajectories and spawning parameters generated for one experiment could be analysed with varying numbers of released gametes. All model results are available in Suppl. S1.

### 2.1 Choice of framework parameters

The range of the framework parameters used for the models was estimated based on literature values, personal communication with foraminifer-researchers, and theoretical mathematical approaches. Five parameters were estimated, and their variation ranges as used in our models are shown in Table 1.

**Table 1.** Summary of framework parameters used for reproduction modelling of planktonic Foraminifera.

|  | Minimum | Maximum |
| --- | --- | --- |
| Experiment duration (h) | 48 | 720 |
| Gamete release synchronization (h) | 12 | 756 |
| Foraminifera density (spec. m<sup>-3</sup>) | 10 | 40 |
| Gamete size (μm) | 2 | 4 |
| Gamete number (*n*) | 12 500 | 400 000 |
| Gamete speed (μm s<sup>-1</sup>) | 25 | 100 |

#### 2.1.1 Synchronization of gamete release

Direct observations of foraminiferal gamete release are rare under laboratory conditions (Bé and Anderson 1976, Spindler et al. 1978, Bé et al. 1983) and have never been observed in nature. Yet, population analyses with high temporal resolution suggested gametogenesis in planktonic Foraminifera to be synchronized in time, with a lunar or semi-lunar cyclicity as a potential trigger for gamete release (e.g. Spindler et al. 1979, Biĵma et al. 1990, Erez et al. 1991, Hemleben and Biĵma 1994, Jonkers et al. 2015, Venancio et al. 2016). Accordingly, for most experiments we assumed a synchronized gamete release within 12 h. However, to explicitly test the effect of a largely unsynchronized release, we modelled several scenarios where the gametes were released over a period between 36 and 756 h.

#### 2.1.2 Foraminifera density in the oceans

To estimate a realistic range of Foraminifera densities in the oceans, we used existing data from research cruises, which took standardized water sample volumes with plankton net tows (Kučera et al. 2013, Rebotim et al. 2017, Visbeck et al. 2017, Kučera et al. 2019, Meilland et al. 2019). For our models, we assume that 10^−40^ spec. m^−3^ are a realistic assumption for planktonic foraminiferal densities for most species. We note that some rare species have lower abundances, and that abundances can be much higher during plankton bloom conditions (Storz et al. 2009).

#### 2.1.3 Size of gametes

The size and shape of planktonic foraminiferal gametes was described in detail for *Hastigerina pelagica* as being ‘3–4 μm in diameter’ (Spindler et al. 1978, p. 429). Other sources assume a gamete size of 3–5 μm (Schiebel and Hemleben 2017), but observations are rare and restricted to a few species. For our models, we used two conservative size estimates of 2 or 4 μm gamete size.

#### 2.1.4 Number of produced gametes per foraminifer

As gametogenesis is not commonly observed in culture (Bé and Anderson 1976, Spindler et al. 1978, Bé et al. 1983) and happens very rapidly (with the gametes leaving the adult shell within seconds) it is challenging to estimate the total number of gametes that are produced by one adult cell. A rough estimate for gamete numbers can be found in Bé and Anderson (1976), who estimated that *Trilobatus sacculifer* produces at least 2.8 × 10^5^ gametes, but who state that probably considerably more gametes had been produced as many had already escaped the shell at the time of observation. Accordingly, Schiebel and Hemleben (2017) estimate the average number of gametes produced by planktonic Foraminifera to be approximately 300 000–400 000.

In addition to these estimates, we used data from studies of calcification intensity in planktonic Foraminifera (Lombard et al. 2010, Naik et al. 2011, Marshall et al. 2013, Weinkauf et al. 2013, Weinkauf et al. 2016, see Suppl. S2) to estimate gamete numbers for a range of species. We used size and weight data of individual foraminiferal shells, together with the density of calcite of *ρ* = 2.7102 g cm^−3^ (Mindat.org Management Team 1993–2020), to estimate the volume of cavities in the shell that can be filled with cytoplasm (Suppl. S3). We further assumed that the shell is filled to 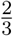 with cytoplasm, and that 30% of the cytoplasm is converted into gametes. The average gamete size was assumed to be 30 μm^3^. Using these parameters, our calculations estimate the production of few tens of thousands to several hundreds of thousands of gametes per individual, depending on the species (Suppl. S4). Convincingly, our estimate for *Trilobatus sacculifer* is *c*.500 000 gametes per individual, which is close to the estimate by Bé and Anderson (1976), and thus supports our calculations. Based on these results, we ran our models “with a range of 12 500 to 400 000 gametes per individual.

#### 2.1.5 Speed of gametes

Gamete speed in planktonic Foraminifera has evaded all investigations so far, and no observations about this topic have been published to our knowledge. However, we used a video by Jennifer Fehrenbacher (Oregon State University, USA) available on YouTube (https://www.youtube.com/watch?v=iCqcKjeqR4g) to estimate gamete speeds. The video shows a specimen of the species *Neogloboquadrina dutertrei* releasing gametes in a culturing dish. We used the size of the foraminifer of 300 μm to approximate the scale of the image, and with this information estimated the distance covered by individual gametes in one second, which appears to be 8–12 gamete body lengths. Accounting for estimation errors, we ran our models with a range of 25–100 μms^−1^ for estimated gamete speeds. The video also shows that gametes are explosively expelled to a distance of *c*.2000 μm, but this initial expulsion is not characteristic for the gametes own motion and was thus not included in our assumptions.

### 2.2 Model description

#### 2.2.1 Module 1—gamete distribution

This module simulated the distribution of gametes by a random walk process with the parameters gamete speed, standard variation of gamete speed, time between direction changes during movement, and maximum survival time of individual gametes. Gamete release was simulated by the placement of 10000 dimensionless particles at the centre of a virtual volume at the beginning of the simulation. The particles moved along a uniformly randomized vector for the given time between direction changes and could not interact with each other. The densities of gametes against the distance from the centre were approximated by a normal distribution for every time step (10 s) of the simulation, up to the defined maximum survival time of the gametes (36 h).

The fit of this approximation was very good, though the approximated gamete densities were biased towards higher concentrations at greater distances as the maximum of the normal distribution showed a skewing to the right (Suppl. S4). Maximal differences (>1%) of modelled and approximated gamete densities always occurred at the initial time steps. For later use in Module 3, the maximum distance travelled against time was approximated by a 2nd-degree polynomial function.

#### 2.2.2 Module 2—Foraminifera trajectories

This module simulated the movement of several adult Foraminifera in a cubic meter of sea water with periodic boundary conditions in a quasi-laminar flow. Starting at random initial positions, Foraminifera were moved with normally distributed randomized speeds (mean: 25 μm s^−1^. standard deviation: 5 μm s^−1^) along a randomly determined fixed starting vector. The movement, vector of individual Foraminifera was altered at random intervals (normally distributed; mean: 600 s, standard deviation: 180 s) by a randomly determined deviation (normally distributed; mean: 0°, standard deviation: 10°, in both *x/y* and *z*-directions). Similar to Module 1, the individual Foraminifera were dimensionless and could not interact.

#### 2.2.3 Module 3—spawning parameter generation

This module generated several values describing the process of spawning or gamete release for the individual Foraminifera. Within the module, all adult cells released their gametes within the synchronization time of the respective experiment. The number of released gametes was generated as a normally distributed random number with a mean according to the settings of the experiment (Table 1) and a standard deviation of 20%. The maximum distance of travel for the gametes was taken from Module 1 results (compare Suppl. S4).

#### 2.2.4 Module 4—determination of gamete encounters

This module determined the probability of encounters of gametes of different adults according to the trajectories determined in Module 2 and the spawning parameters determined in Module 3. For each time step of the simulation, the distances between all individual Foraminifera were compared against the maximum gamete travel range of involved adults (from Module 1) and the status of spawning (time between gamete release and maximum life expectancy of gametes). The list of probable gamete interaction events was stored for final analysis in Module 5.

#### 2.2.5 Module 5—calculation of gamete fusions

This module determined the number of gamete fusions into zygotes from the list of probable gamete interaction events produced by Module 4. Since our simulated gametes perform a random walk, we used the equation for particle collisions from kinetic gas theory (Eq. 1).

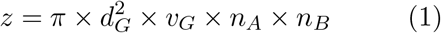

where *z* is the collision frequency, *d_G_* is the gamete diameter, *v_G_* is the gamete velocity, and *n_A_* and *n_B_* are the concentrations of the gametes of Foraminifera individuals *A* and *B*.

The validity of this approximation was tested by comparing the collision rates obtained from Eq. (1) with direct numerical simulations of the upper triangular matrix of gamete concentrations in the set [5 … 100, 5] gametes ml^−3^. This comparison was replicated for all gamete speeds used in the different experiments and is summarized in Suppl. S4. Root mean square errors were generally larger for low gamete concentrations, suggesting that the differences between the approaches are due to sampling size rather than being systematic.

### 2.3 Data analysis

All model results have been analysed in R v. 4.0.2 (R Core Team 2020). The relationship between model parameters and reproductive success was modelled using generalized additive models (GAMs) of the form presented in Eq. (2) as implemented in the R-package ‘gamlss’ v. 5.1-7 (Rigby and Stasinopoulos 2005), using their own fitting algorithm. Dependent variables in the GAMs were fitted via maximum likelihood with P-splines based on singular value decomposition. The family of the link-function for the GAMs was chosen based on an evaluation of the distribution of the dependent variable in the R-package ‘fitdistrplus’ v. 1.1-1 (Delignette-Muller and Dutang 2015).

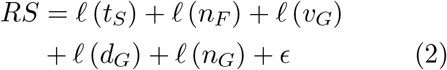

where *RS* is the reproductive success parameter to be modelled, *t_S_* is the degree of synchronization (i.e. time-window size during which gametes are released), *nf* is the density of Foraminifera, *v_G_* is the gamete velocity, is the gamete diameter, *n_G_* is the number of gametes released per foraminifer, *ϵ* is the error term, and *ℓ* denotes a P-spline smooth.

The variation of reproductive success parameters was calculated as coefficient of variation 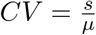, where *s* is the standard deviation and *μ* is the mean of the parameter. To estimate the distribution pattern of reproductive success across individuals (i.e. whether all individuals had the same reproductive success), we used Ripley’s *L* function (Ripley 1976) and visualized influential parameters across the space of largest variation using flexible discriminant analysis (FDA; Hastie et al. 1994) as implemented in the R-package ‘mda’ v. 0.5-2. The survival rate of planktonic Foraminifera has been estimated using non-linear regression.

## 3 Results

The results of our models suggest that different experimental setups strongly vary in their individual reproductive success. Between 0 and 39 (mean: 5.3, 3rd quartile: 5) individual Foraminifera were able to successfully reproduce per experimental run, generating a total of 0 to 228 000 (mean: 604.3, 3rd quartile: 16.35) zygotes. With these data, we evaluated reproductive success in two ways: (1) We estimated the number of zygotes produced (i.e. successful fusions of two gametes) per successful individual and in relation to the population. These numbers are critical, as sufficient zygotes are required to sustain the population. (2) We estimated the proportion of the population that was able to successfully reproduce and the distribution of reproductive success across individuals. This is a quantity for the reproductive success amongst planktonic Foraminifera, which has an impact on population viability and gene flow.

### 3.1 Scale of reproductive success

We find that across all experiments, between 0 and 114000 zygotes are produced by each successful individual foraminifer; with a mean of 94 and a 3rd quartile of 2. Across the entire population, compensating for unsuccessful individuals, this translates to 0–11400 zygotes per foraminifer (mean: 21, 3rd quartile: 1).

The individual success in producing zygotes follows a strongly positively skewed gamma distribution, containing large numbers of zero-values. We therefore chose a zero-adjusted gamma distribution as link-function for the GAM (results are shown in Table 2). We And that when only considering the successful individuals, synchronization time and Foraminifera density do not significantly influence reproductive success. Gamete speed shows a negative relationship with zygote production, while both gamete size and the number of gametes produced per foraminifer positively influence the number of fusions (Suppl. S4). The coefficient of variation of fusions per individual ranges between 0.78 and 20.37, with a mean at 4.50 and 3rd quartile at 5.07. The variation follows a gamma distribution, and the fitted GAM implies that variation increases with a relaxation of synchronization and decreases with foraminiferal density and gamete speed (Suppl. S4). Conversely, gamete size and number of gametes produced do not affect the variation of reproductive success.

**Table 2.**
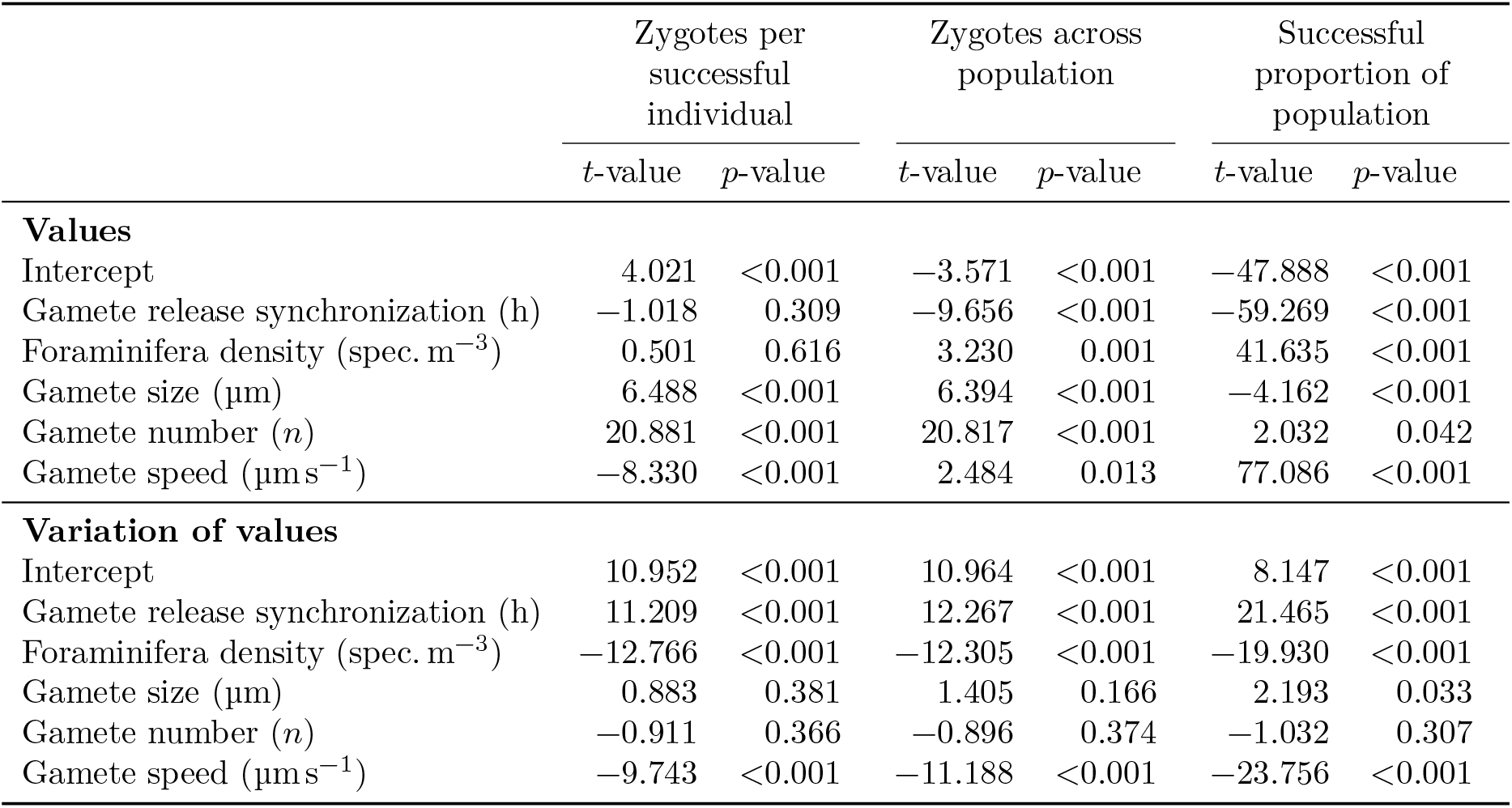
Results from a generalized additive model of foraminiferal reproductive success in dependence of model parameters.

The interpretation changes, however, when considering zygote production across the entire population (i.e. including unsuccessful individuals), which is more relevant for the retention of the population (Table 2, Fig. 1a–b). All modelled parameters significantly influence reproductive success, with the success rate increasing with stricter synchronization of gamete release and increases in foraminiferal density, and number, size, and speed of gametes. The coefficient of variation of zygote production across the population ranges between 0.77 and 20.37, with a mean of 4.35 and a 3rd quartile of 4.88. The variation of reproductive success increases with a relaxation of the synchronization of gamete release but can be decreased by higher population densities and greater gamete speed, while size and number of gametes exhibit no controls on the variation of reproductive success across the population (Fig. 1c).

**Figure 1.**
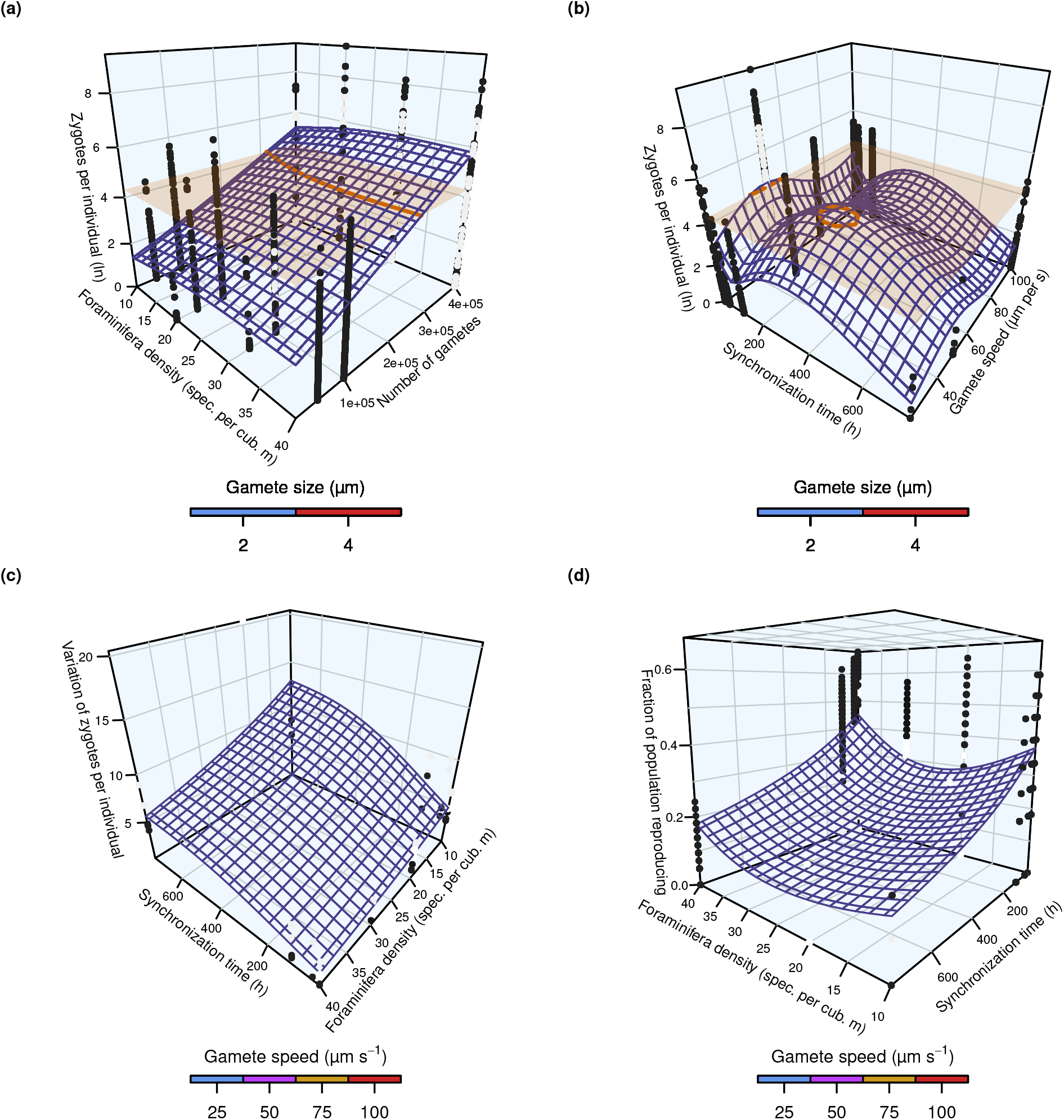
Reproductive success of planktonic Foraminifera depending on model parameters based on a generalized additive model. (a–b) Number of zygotes produced per specimen across the population (log_e_-transformed). Orange horizontal plane and trace indicate 75 zygotes per foraminifer. (c) Coefficient of variation of zygotes per individual across population. (d) Fraction of the population successfully reproducing depending on model parameters.

### 3.2 Proportion of the population reproducing

The reproductive success of the population is expressed as the fraction of individuals that were participating in at least one fusion of two gametes into a zygote. Within our model framework, between 0 and 97.5% of the population reproduced successfully, with a mean of 17.7% (3rd quartile: 26.7%).

The population’s reproductive success follows a zero-inflated beta distribution. The GAM indicates that the reproductive success is influenced by all tested parameters: Stronger synchronization, higher foraminifer densities, and smaller, faster and more numerous gametes positively influence reproductive success (Table 2, Fig. 1d, Suppl. S4). The coefficient of variation of the population-wide success rate ranges between 0.07 and 10.00, with a mean of 1.54 (3rd quartile: 1.83). The GAM on the gamma distribution shows an increase of variation with relaxed synchronization, lower foraminiferal density, and larger, slower gametes; the number of gametes exhibits no effect on success variation (Table 2, Suppl. S4).

However, even in a scenario where a large proportion of the population reproduces successfully, it is possible that the majority of zygotes is produced by only a few specimens. This can be tested by looking at the distribution of two parameters: (1) The number of successful reproductions of each individual with another foraminifer and (2) the proportion of zygotes produced by each individual. Ripley’s *L* function implies that the number of successful reproductions with different partners falls in either of two groups across our experiments (Fig. 2a, c): (1) Some experiments show a low divergence following a shallow beta distribution or coming close to a normal distribution (Suppl. S4). Here, many individuals reproduce with a moderate number of different partners, but relatively few are either unsuccessful or reproduce with extensive numbers of other individuals. Around half of the experiments (52%) fall in this group. (2) In other experiments, the divergence follows a steep beta distribution, with most of the population reproducing with none or very few other individuals, while few individuals perform excessively well. An FDA shows that higher gamete production and population density promote low divergence, while a relaxed synchronization time causes high divergence in the number of successful reproductions, For the number of produced zygotes per individual, all experiments follow a relatively steep beta distribution, but the divergence is lower in some experiments than in others (Fig. 2b, d, Suppl. S4). The low-divergence group (13% of all experiments) shows large numbers of individuals that do not produce any zygotes at all, but a rather uniform distribution of all successful individuals between very few and numerous zygotes produced. Within the high-divergence group, few individuals produce the vast majority of zygotes. A more even distribution is promoted by larger, faster gametes and a higher population density, while relaxed synchronization and a higher number of gametes bolster a high divergence of the number of produced zygotes.

**Figure 2.**
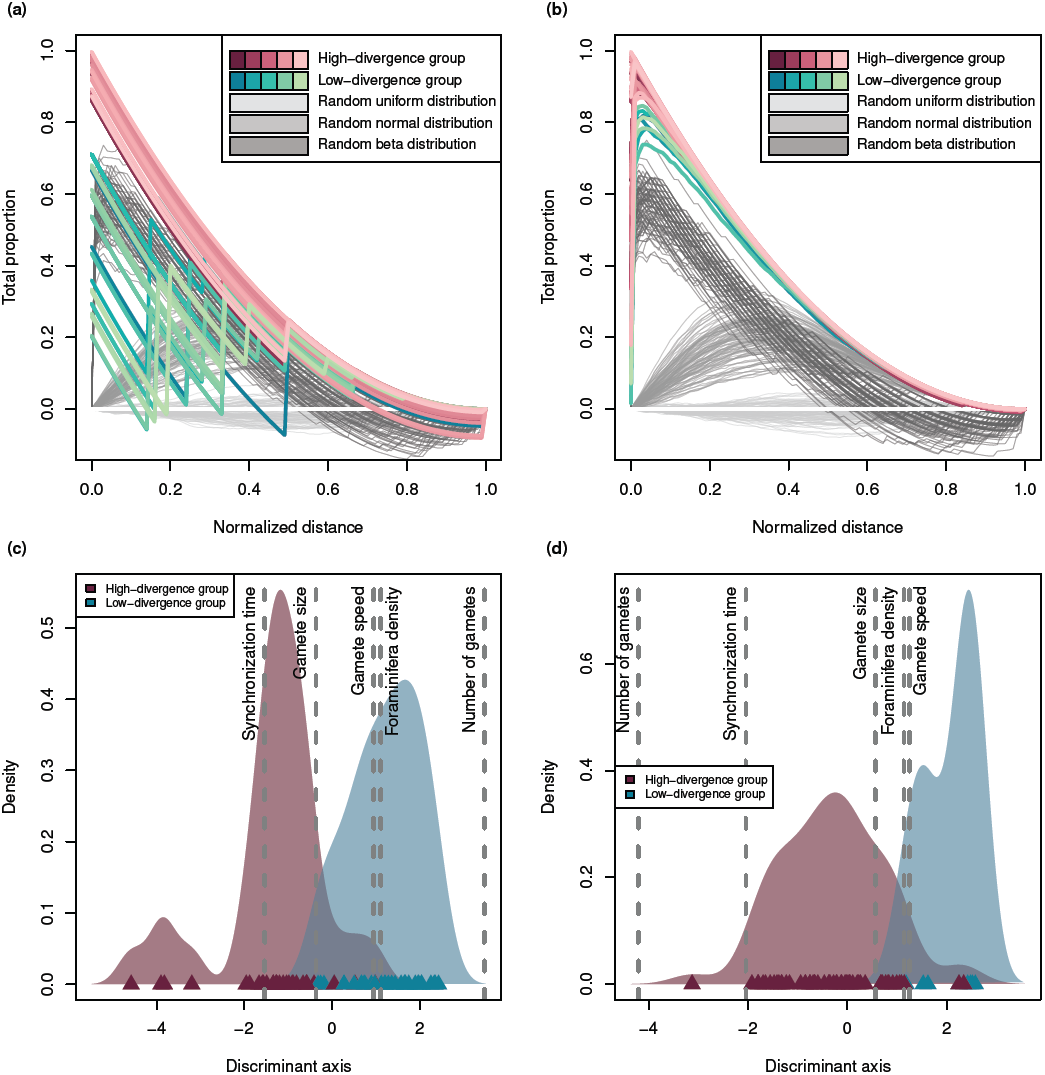
Divergence of reproductive success across individuals in planktonic Foraminifera under varying model setups. (a–b) Ripley’s *L* functions of number of reproductive events with another individual (a) and total number of zygotes produced per individual (b). The experiments were categorized into groups with higher and lower divergence between individuals, respectively: Under low divergence, all individuals have comparable success, under high divergence some individuals are very successful at the expense of others. Random examples (100 replications) of a uniform, normal, and beta distribution are shown for comparison. (c–d) Flexible discriminant analysis of the experiments, categorized according to Ripley’s *L* functions, for number of reproductive partners (c) and number of zygotes produced (d). Data points are shown as triangles, with kernel density function as shaded area. Influence direction and weighting of different experimental parameters is indicated by grey, dashed lines.

## 4 Discussion

### 4.1 Individual reproductive success and the sustainability of the population

To evaluate the reproductive success necessary for maintaining a population, data for the survivability of zygotes in planktonic Foraminifera are needed. The survivability of the larvae of Metazoa has been studied to some extent (Dahlberg 1979, Anderson 1988, Rice et al. 1993, Graham et al. 2008), but no such data are available for planktonic protists. We thus estimated the probability of survival of planktonic foraminiferal zygotes to the reproductive stage. Brummer et al. (1988b, fig. 3) provide data on planktonic foraminiferal shell sizes across all species down to 60 μm. The size-range over which representative abundance data had been collected in that study allowed us to fit an exponential model that is indicative of the survival of planktonic Foraminifera during their ontogeny (Fig. 3). Brummer et al. (1988a) estimate a proloculus size (the initial shell of the foraminifer, consisting only of one chamber) of 12–20 μm. We thus assumed a zygote size of 15 μm and consider a foraminifer with a shell size of 125–150 μm to be of reproductive age (Peeters et al. 1999). Using our survival curve, we could estimate the survival rate of planktonic Foraminifera from the zygote to the reproductive stage as approximately 5% (Fig. 3). This aligns with comparable estimates for benthic Foraminifera (Suppl. S4) and corresponds to observed survival curves in multicellular organisms (Dahlberg 1979, Rice et al. 1993). Under this assumption, an average reproductive success of >70–100 zygotes per individual foraminifer would be sufficient to sustain a viable population.

**Figure 3.**
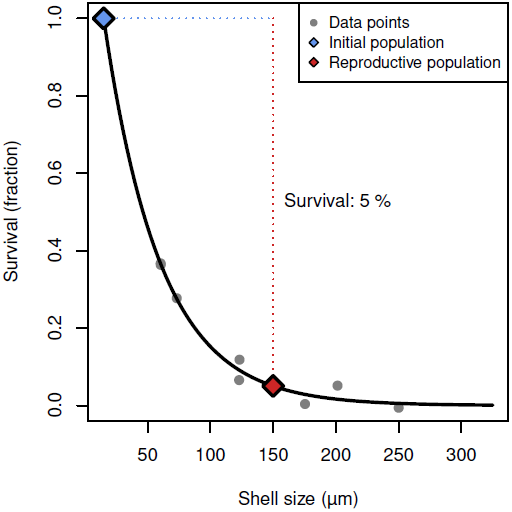
Estimated survival rate of planktonic Foraminifera from the zygote (initial population, 15 μm) to the reproductive stage (150 μm), based on plankton net and sediment data from Brummer et al. (1988b).

However, only 9 % of our experiments reach these numbers (compare Suppl. S1) and they all have in common a stringent synchronization of gamete release within 12 h, the production of 400 000 gametes per specimen, and population densities of ≥20 spec. m^−3^. We further note, that all models that could sustain the population show relatively low gamete speeds of *c*.50μm s^−1^. This corresponds to the region of successful reproduction shown in Fig. 1a, b (above the orange threshold plane) which indicates large numbers of gametes, high population densities, and strict synchronization as prerequisites.

#### 4.1.1 Temporal and spatial synchronization of reproduction

Planktonic Foraminifera generally occur in very low abundances of only a few to some tens of specimens per cubic metre in the world’s oceans (e.g. Kučera et al. 2013, Meilland et al. 2019). Some kind of temporal synchronization of their reproductive cycles was therefore suggested as requirement to sustain a population, driven by either lunar cyclicity (Spindler et al. 1979, Bijma et al. 1990, Erez et al. 1991, Jonkers et al. 2015, Jentzen et al. 2019) or alternative higher-order cyclicities (Schiebel and Hemleben 2005, Jonkers and Kučera 2015). Our data support the need for a temporally synchronized reproduction, because even under ideal circumstances a lack of temporal synchronization leads to unsustainably low numbers of gamete fusions (Suppl. S1). Additionally, a stronger synchronization reduces the variation in individual reproductive success (Fig. 1c), allowing stable population sizes to be maintained more easily. Triggers for gamete generation and their synchronized release may include lunar tides or other physical oceanic parameters such as turbulence, turbidity, or temperature (Norris 2000).

Temporally synchronized reproduction could be further aided by a spatial concentration of the reproductive efforts, for instance by sinking of reproductive specimens to a physical boundary layer like the halocline. Such a cyclic depth migration has been suggested for planktonic Foraminifera (Erez and Honjo 1981, Eggins et al. 2003, Schiebel and Hemleben 2005) and also other marine microplankton (Font-Muñoz et al. 2019). The turbulent nature of the ocean seems to promote the concentration of plankton into spatial clusters and even pairing of specimens for reproductive purposes (Borgnino et al. 2019, Basterretxea et al. 2020). Based on our findings, and in alignment with empirical observations (Meilland et al. 2019), we thus speculate that local population densities must be increased through some kind of spatial concentration to reach the necessary population densities of ≥20 spec. m^−3^ implied by our model. Our models used a laminar flow to reduce the complexity and associated calculation times, so we may have underestimated the effective population densities, but certainly not drastically. Consequently, spatial concentration combined with temporal synchronization of planktonic Foraminifera seems to be the only way to sustain populations in a natural environment under the assumption of purely sexual reproduction (compare Fig. 1a, b). The spatial concentration-effect could be aided further if planktonic Foraminifera were to adopt a pseudo-benthic lifestyle for the time of reproduction by attaching to marine snow, as was speculated by Hilbrecht and Thierstein (1996). Further, chemical signalling between planktonic Foraminifera after concentration/pairing, which could trigger gamete production and has been observed in diatoms (Basu et al. 2017), could farther enhance synchronization.

#### 4.1.2 Gamete production and characteristics

We observe that larger size and speed of gametes positively affects reproductive success across the population (Fig. 1b). This is likely because faster gametes can more easily reach the nearest gamete cloud of another individual and larger gametes have a higher random chance of collision. Interestingly, when only considering successful individuals, a lower gamete speed is beneficial for reproductive success (Suppl. S4). We speculate that in this case, the gamete clouds of two individuals are already overlapping, and slower gametes maximize the local concentration of gametes due to reduced spreading. It thus seems that gamete speed is governed by a delicate trade-off to maximize reproductive success. These assumptions could be somehow relaxed if foraminiferal gametes had a form of active partner-detection mechanism. For instance, for some algae, a phototactic behaviour has been described (Togashi et al. 1999), which would ensure a uniform direction of movement of gametes to increase encounter rates. Chemotaxis could increase gamete encounter rates even more by enabling active partner tracking, if foraminiferal gametes overcame the sensitivity limit by, for instance, having sufficient receptors on their cell surface (Bialek and Setayeshgar 2005). We decided against including such mechanisms in our models for lack of evidence but suggest concentrating empirical scientific research in this area in the future.

Our models further suggest that populations would not be sustainable in species producing less than *c*.250 000 gametes (Fig. 1a). In our estimates of gamete production rates, only the exceptionally large species *Trilobatus sacculifer* and *Orbulina universa* generate that many gametes. All other species, thus, would not be able to maintain their populations. We note, that we may have underestimated the number of gametes for some species, but for *T. sacculifer* our calculation matches well with the numbers reported by Bé and Anderson (1976). We argue that one possible solution is that smaller species produce a greater number of smaller gametes as opposed to the fewer, larger gametes in larger species. In this case, the smaller gametes would likely have fewer energy reserves in the form of lipid droplets (Bé et al. 1983), reducing their survival time. However, this could be offset by the increased encounter rates generated by the increased gamete counts. Should this hypothesis be true, the equivalent of *r*- and *K*-strategists may exist amongst planktonic Foraminifera, with potential implications for the adaptability of different species. Alternatively, if a strong spatial concentration or pairing of individuals took place prior to gamete release, this could levy the issue of low gamete counts by significantly raising encounter rates.

We argue that our results make the existence of mating types, as was suggested for other protists (Power 1976), unlikely in planktonic Foraminifera. Their existence would further reduce encounter rates between suitable gametes, making a viable population nearly impossibleto achieve. This also corresponds with the observation that planktonic foraminiferal gametes are isogamous (Goldstein 1997), while mating types overwhelmingly induce anisogamy.

Due to the difficulty of long-term culturing of planktonic Foraminifera, little is known about the nature of the gametes. For example, it has never been investigated if the released gametes are indeed haploid gametes, or rather diploid swarmers that would grow to an adult foraminifer without the need to fuse with another swarmer. Should this be the case, it would alleviate the problem of managing to ensure encounters, and genome sequencing of adult Foraminifera and their gametes may be able to resolve this issue in the Future. Nevertheless, from scarce laboratory observations, it seems that the released gametes do not grow into adult foraminifers, and that asexual reproduction occurs rarely in planktonic Foraminifera (Takagi et al. 2020). We have no evidence for the existence of persistent asexual reproductive phases in planktonic Foraminifera as they are known for benthic Foraminifera, but it may be a way to reduce the necessary success rates during the sexual reproductive phases as we modelled them here.

### 4.2 Distribution of reproductive success across the population

Interestingly, planktonic Foraminifera are the only major group of marine protists believed to reproduce purely sexually through gametes. While the majority of protists is capable of sexual reproduction, usually they reproduce asexually, which is frequently the dominant mode of reproduction (Archibald et al. 2017). This is even more intriguing when considering that planktonic Foraminifera descended from benthic Foraminifera, which are well known for having both sexual and asexual reproductive cycles (Goldstein 2002). It appears plausible that planktonic Foraminifera secondarily reduced their asexual reproductive cycle (Kučera et al. 2017), considering that sexual reproduction is the more ancient reproductive mode (Goldstein 1997, Speijer et al. 2015). This assumption, however, raises two main questions: (1) If asexual reproduction was secondarily reduced in all planktonic Foraminifera, this would have had to occur at least twice independently, since evidence suggests that modern planktonic Foraminifera are polyphyletic (Darling et al. 1997). (2) Sexual reproduction conveys several benefits, like the generation and spread of beneficial mutations (Kondrashov 1988, Speijer et al. 2015) and the purging of harmful mutations (Muller 1964). Nevertheless, sexual reproduction always bears a risk of failure, since it depends on the encounter of the reproductive cells of two individuals. Chances for success are decreased in organisms that (a) simply release their reproductive cells instead of engaging in any form of pairing and (b) occur in small abundances. Both is ostensibly true for planktonic Foraminifera, and they would be an oddity in the protist world if they truly relied solely on sexual reproduction. We, thus, suggest further studies in this field to gain more insights into the matter and evaluate if not asexual reproduction plays a more important role in planktonic Foraminifera than is currently believed (compare Davis et al. 2020, Takagi et al. 2020).

Interestingly, the models that would allow the largest fraction of the population (>40%) to reproduce successfully are decidedly not the models that allow sustainable numbers of successful gamete fusions (Suppl. S1). The production of medium numbers of gametes (50 000–100 000) with higher speeds (75–100 μm s^−1^) seem to be most beneficial for an even divergence of success across the population (Fig. 1d, Suppl. S4). In contrast, the six models which lead to sustainable populations, have only 25% of the population successfully reproducing on average. It seems as if the framework parameters which benefit population-preservation the most are rather detrimental for the upkeep of the gene flow and genetic intermixing within the population. The only exception is a strict temporal synchronization of the reproduction, which reduces variation in the population-wide success and thus increases the odds for an even genetic inheritance (Suppl. S4). This discrepancy does not seem to stem from the distribution of the number of reproductive events across individuals, as shown by Ripley’s *L* (Fig. 2a, Suppl. S4). All models that sustain the population show a low divergence of reproductive events, meaning that the majority of the population reproduces with a medium number of other individuals, thus keeping gene flow on a moderate level. Rather, it is the number of zygotes produced during these gamete cloud encounters, that is skewed toward a large discrepancy between individuals. Each successful encounter of gamete clouds between two individuals can produce between one and hundreds of thousands of zygotes, depending on how many gametes of the tow adult cells manage to fuse. In all models that can sustain the population, only few gamete cloud encounters produce large numbers of zygotes, so that the majority of zygotes in the population is produced by the minority of Foraminifera (Fig. 2b, Suppl. S4). This corresponds with the observation that especially the high number of gametes necessary to sustain the population in our models increases the divergence in the number of zygotes individual Foraminifera produce (Fig. 2d).

We could assume that producing sufficient numbers of zygotes is more important to species survival than an even distribution of reproductive success across all individuals, although a combination of both would arguably be most beneficial. Accordingly, framework parameters maximizing zygote production would be favoured in terms of evolutionary fitness, even if it means sacrificing population-wide reproductive success and, thus, gene flow. Tf this held true, it could help explaining some prior observations: (1) Norris (2000) discussed in detail the problem of diversification and speciation in the open ocean plankton, which lives in an environment void of physical barriers which could prevent gene flow. Should planktonic Foraminifera indeed sacrifice gene flow for successful reproduction on the population level, it may help explain speciation between genetically isolated populations, even if they occur in sympatry (de Vargas et al. 1999, Morard et al. 2011, Weiner et al. 2012) or undergo speciation in a homogeneous environment (Kučera et al. 2017;. (2) It could be shown that planktonic Foraminifera communities, when exposed to stressful environments, most often react with an adaptive response based on pre-existing variability rather than innovation through evolution (Weinkauf et al. 2014, Brombacher et al. 2017, Weinkauf et al. 2019). The evolvability of planktonic Foraminifera may just be inherently low (compare West-Eberhard 2003, Wagner 2011), or alternatively it may be that innovations simply cannot be fixated in the population due to reduced gene flow, which could in turn explain the spikes in variation sometimes observed in fringe environments (Weinkauf et al. 2014).

## Acknowledgements

We thank Jelle Bijma (Alfred-Wegener-Institut, Helmholtz-Zentrum fÃ(Er Polar- und Meeres-forschung, Bremerhaven, Germany), Geert-Jan Brummer (Koninklijk Nederlands Instituut voor Onderzoek der Zee, Texel, The Netherlands), Susan T. Goldstein (University of Georgia, Athens, USA), Katsunori Kimoto (Japan Agency for Marine Earth Science and Technology, Yokosuka, Japan), Michal Kučera (Zentrum für Marine Umweltwissenschaften, Bremen, Germany), and Howard J. Spero (University of California—Davis, Davis, USA) for fruitful discussions of framework parameters and foraminiferal reproduction in general. Patrick Gonzalez (Université de Genève. Geneva, Switzerland) is thanked for providing the hardware to run some of the models. MFGW was partly financed through PRIMUS and PROGRES Q45 grants during the work on this study.

## CRediT author contributions

|  |  |
| --- | --- |
| Conceptualization: | MFGW, MS, AKMW |
| Methodology: | MFGW, MS, AKMW |
| Software: | MS |
| Validation: | MS, AKMW |
| Formal analysis: | MFGW |
| Investigation: | MFGW, MS, AKMW |
| Resources: | MS |
| Data curation: | MFGW, MS |
| Writing – original draft preparation: | MFGW, MS, AKMW |
| Writing – review and editing: | MFGW, MS, AKMW |
| Visualization: | MFGW |
| Funding acquisition: | MFGW |

