## Supplement S3 for "Reproduction of a marine planktonic protist: Individual success versus population survival"

#Estimate biovolume of protist shells based in shell weight and size
#Author: Manuel F. G. Weinkauf
#Version: 1.0
#Date: 7 February 2020

#+++++++++++++++++++++++++++++++++++++++++++++++++++++++++++++++++++++++++++++++++++++++++++++++++++++++++++#
#This work is licensed under the Creative Commons Attribution-NonCommercial-ShareAlike 3.0 Unported License.#
#To view a copy of this license, visit http://creativecommons.org/licenses/by-nc-sa/3.0/.                   #
#+++++++++++++++++++++++++++++++++++++++++++++++++++++++++++++++++++++++++++++++++++++++++++++++++++++++++++#

#########################################################################
# Function to calculate biovolume of protists                           #
# Necessary input variables:                                            #
#    Size: Size of the shells as cross-sectional area in square         #
#          micrometre.                                                  #
#          *numeric*                                                    #
#    Weight: Weight of the shells in microgram.                         #
#            *numeric*                                                  #
#    Type: Geometric type of the shell.                                 #
#          *character*, one of "spherical", "globorotalid",...          #
#          "cylindrical", or "truncatulinoid"                           #
#    cut.off: What portion of the shell volume is filled with...        #
#             cytoplasm (fraction).                                     #
#             *numeric*                                                 #
#             default=2/3                                               #
#    to.gametes: What proportion of the cytoplasm is transferred into...#
#                the gametes (fraction).                                #
#                *numeric*                                              #
#                default=0.3                                            #
#    gamete.volume: Volume of one gamete in cubic micrometre.           #
#                   *numeric*                                           #
#                   default=30                                          #
#    Density: Density of shell material (in g/cubic cm).                #
#             *numeric*                                                 #
#             default: 2.7102                                           #
#    Samples: Sample names for results table.                           #
#             *character*                                               #
#    Species: Species name.                                             #
#             *character*                                               #
# Output data: Number of gametes produced by protists.                  #
# Input dataset: Sizes and weights of prostist shells.                  #
#########################################################################

Biovol<-function(Size, Weight, Type, cut.off=2/3, to.gametes=0.3, gamete.volume=30, Density=2.7102, Samples, Species) {
	#Test data consistency
	if (length(Size)!=length(Weight)) {stop("Input parameters of unequal length!")}
	if (length(Samples)!=length(Size)) {stop("Samples must be of same length as data!")}
	if (length(Species)!=length(Size)) {stop("Species must be of same length as data!")}
	if (!all(Type %in% c("spherical", "globorotalid", "cylindrical", "truncatulinoid"))) {stop("Type must be one of 'spherical', 'globorotalid', 'truncatulinoid', or 'cylindrical'")}
	{if (length(Type)==1) {
		Type<-rep(Type, length(Size))
		warning("Type has length one and will be assumed to be true for all specimens.")
		}
	else if (length(Size)!=length(Type)) {stop("Type must be of either length one or same lenght as other input parameters.")}
	}
	
	#Set up results table
	Res<-as.data.frame(matrix(NA, length(Size), 8))
	colnames(Res)<-c("Sample", "Species", "Volume.cell", "Volume.calcite", "Volume.cavity", "Volume.cytoplasm", "Volume.gametes", "Number.gametes")
	Res[,"Sample"]<-Samples
	Res[,"Species"]<-Species
	
	#Scale size as volume
	for (i in 1:length(Size)) {
		{if (Type[i]=="spherical") {Res[i,"Volume.cell"]<-(4/3)*pi*(sqrt(Size[i]/pi))^3}#Sphere
		else if (Type[i]=="globorotalid") {Res[i,"Volume.cell"]<-2*((((sqrt(Size[i]/pi)/2)*pi)/6)*(3*sqrt(Size[i]/pi)^2+(sqrt(Size[i]/pi)/2)^2))}#Two conjoined sphere segments
		else if (Type[i]=="cylindrical") {Res[i,"Volume.cell"]<-pi*sqrt(Size[i]/pi)^2*sqrt(Size[i]/pi)}#cylinder
		else if (Type[i]=="truncatulinoid") {Res[i,"Volume.cell"]<-(1/3)*pi*sqrt(Size[i]/pi)^2*sqrt(Size[i]/pi)}#cone
		}
	}
	
	#Scale density as microgram/cumum
	Density<-(Density*1000000)/1000000000000
	
	#Calculate amount of calcite in shell
	Res[,"Volume.calcite"]<-Weight/Density
	
	#Calculate bio-volume
	Res[,"Volume.cavity"]<-Res[,"Volume.cell"]-Res[,"Volume.calcite"]
	Res[,"Volume.cytoplasm"]<-Res[,"Volume.cavity"]*cut.off
	Res[,"Volume.gametes"]<-Res[,"Volume.cytoplasm"]*to.gametes
	Res[,"Number.gametes"]<-Res[,"Volume.gametes"]/gamete.volume
	
	#Return results
	return(Res)
}
