## Supplement S4 for "Reproduction of a marine planktonic protist: Individual success versus population survival"

bioRxiv

*Manuel F. G. Weinkauf*[1,2,3], *Michael Siccha*[1], *and Agnes K. M. Weiner*[1]

[1]Center for Marine Environmental Sciences (MARUM), Universität Bremen, Leobener Str. 8, 28359 Bremen, Germany; [2]Department of Earth Sciences, Université de Genève, Rue des Maraîchers 13, 1205 Genève, Switzerland; [3]Institute of Geology and Palaeontology, Univerzita Karlova, Albertov 2038/6, 128 43 Praha, Czech Republic

### 1 Gamete production and modelling

**Table S1.** Estimated gamete production of different species of planktonic Foraminifera based on shell size and weight data (compare Fig. S1 and Suppl. S1).

| Species | Median ($n$) | Std. error ($n$) |
|---|---|---|
| *Globigerina bulloides* | 32 635 | 1987.46 |
| *Globigerinoides elongatus* | 40 751 | 2029.05 |
| *Globigerinoides ruber ruber* | 38 233.5 | 22 066.71 |
| *Globigerinoides ruber albus* | 19 620 | 1752.75 |
| *Globorotalia inflata* | 21 982.5 | 438.35 |
| *Globorotalia scitula* | 12 196 | 266.74 |
| *Orbulina universa* | 286 958 | 14 362.51 |
| *Trilobatus sacculifer* | 494 478 | 30 472.56 |

**Table S2.** Summary information of Modules 1 and 3 of the reproduction model. Bias and root-mean-square errors (RMSE) of Module 1 of the gamete distribution model are low. The coefficient of determination for the 2nd-degree polynomial, with which the maximal gamete range for Module 3 was estimated, shows very good results.

| | Module 1 | | Module 3 |
|---|---|---|---|
| Gamete speed (μm $s^{-1}$) | Bias (%) DNM-model | RMSE (%) | $R^2$ |
| 25 | −0.0045 | 0.300 | 0.9732 |
| 50 | −0.0021 | 0.150 | 0.9633 |
| 75 | −0.0014 | 0.100 | 0.9705 |
| 100 | −0.0011 | 0.077 | 0.9762 |

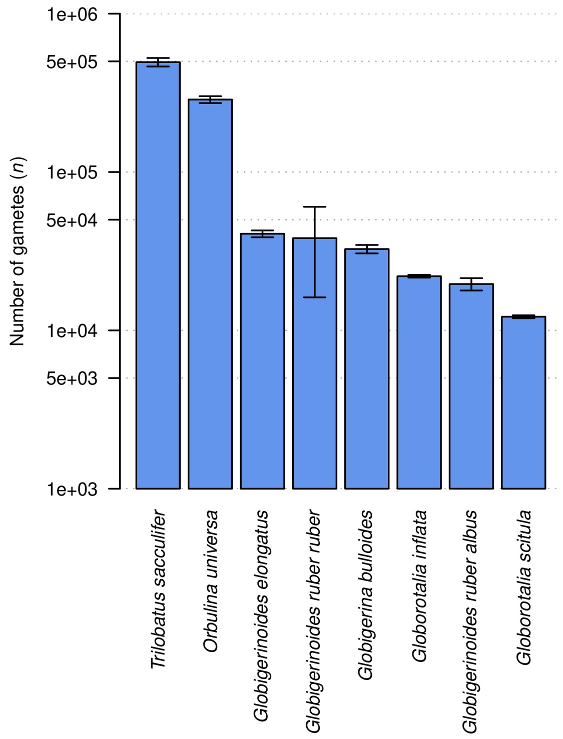


**Fig. S1.** Estimated number of produced gametes across different species of planktonic Foraminifera, based on size–weight data (compare Table S1). Error bars indicate standard error of the estimate.

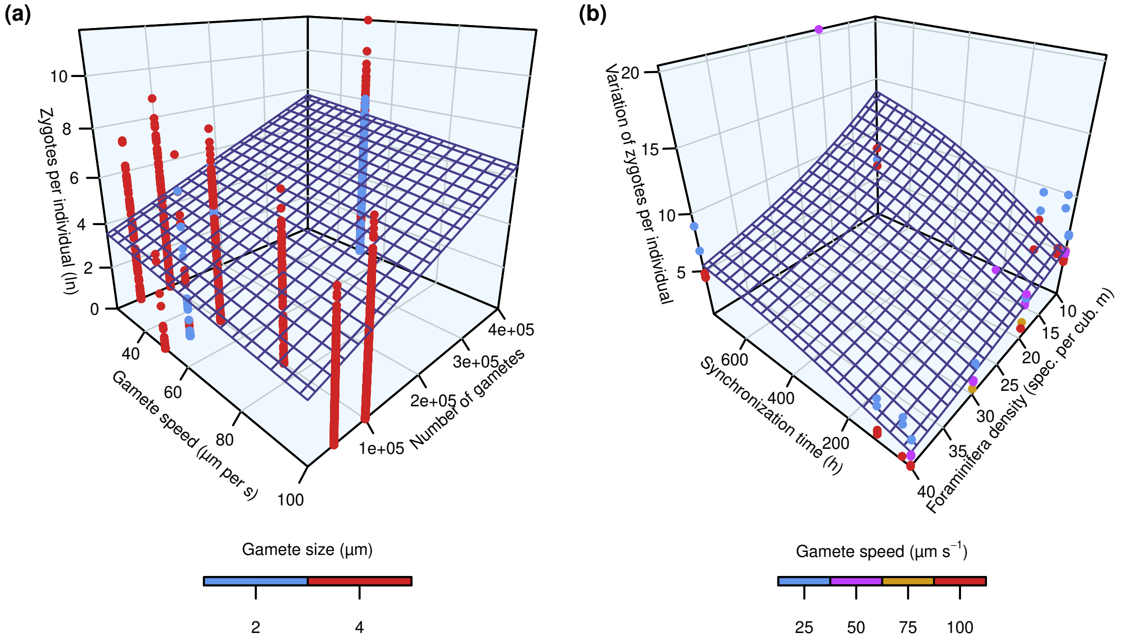


**Fig. S2.** Reproductive success of planktonic Foraminifera depending on model parameters based on a generalized additive model. (a) Number of zygotes produced per successful specimen ($\log_e$-transformed). (b) Coefficient of variation of zygotes per successful individual.

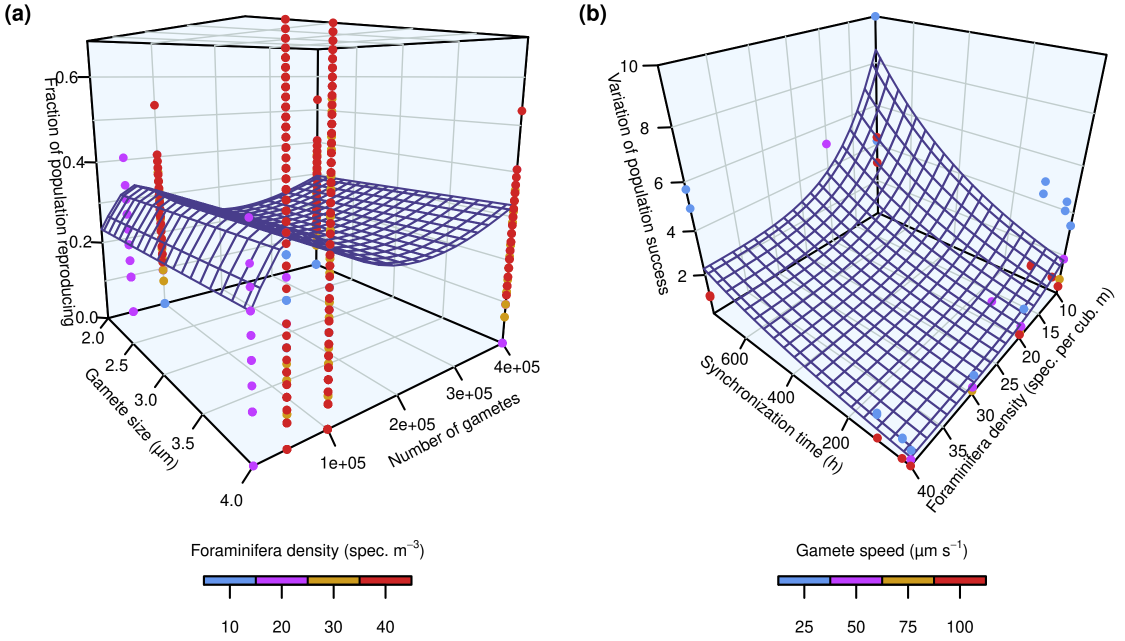


**Fig. S3.** Reproductive success of planktonic Foraminifera depending on model parameters based on a generalized additive model. (a) Fraction of the population successfully reproducing depending on different model parameters. (b) Coefficient of variation of fraction of the population successfully reproducing.

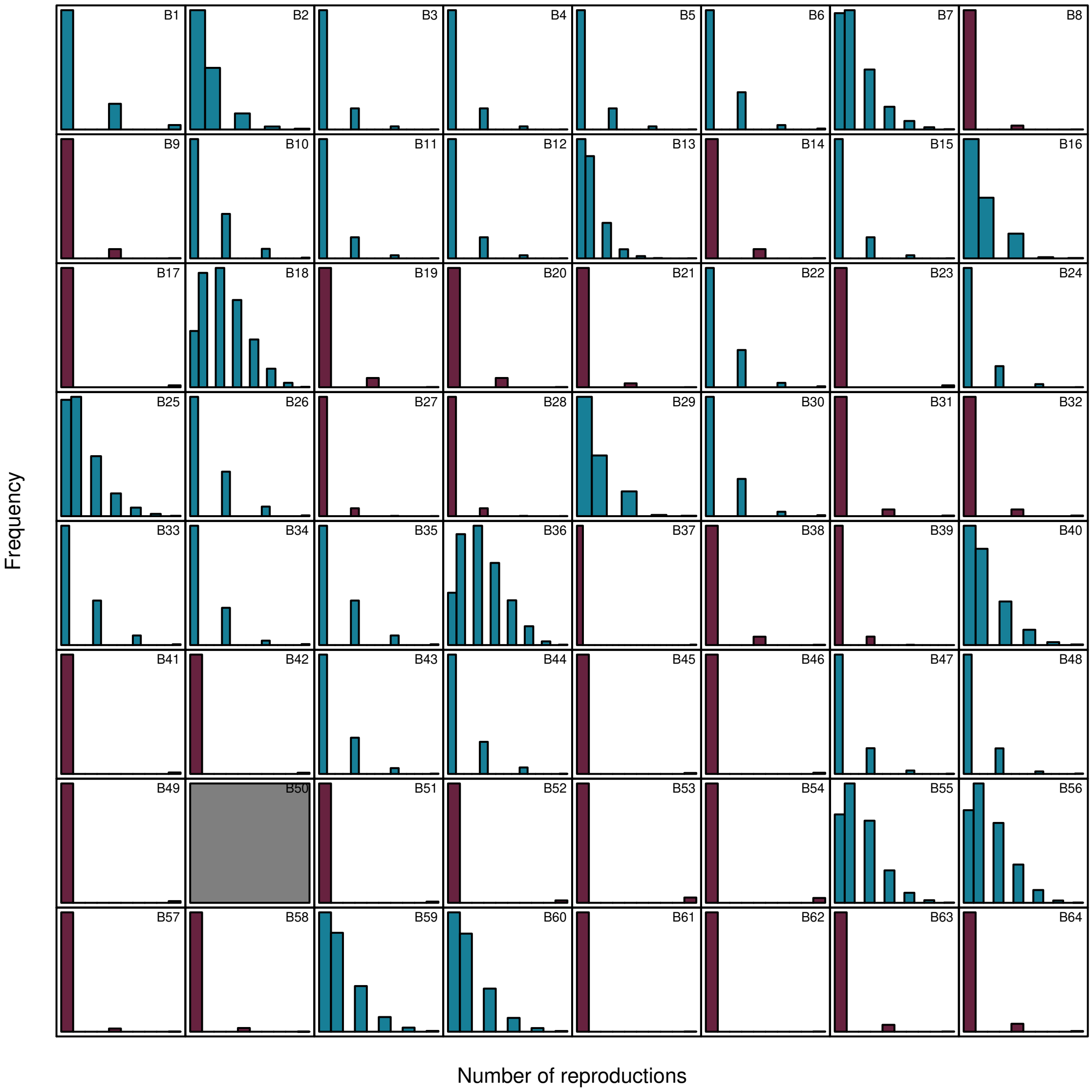


**Fig. S4.** Histograms of the number of reproductive successes (i.e. at least one fusion of gametes with a distinct partner) per individual planktonic foraminifer across all experiments. Experiments resulting in low dispersion shown in teal, those with high dispersion shown in burgundy (compare article for details). Experiment B 50 showed no successful reproductive events.

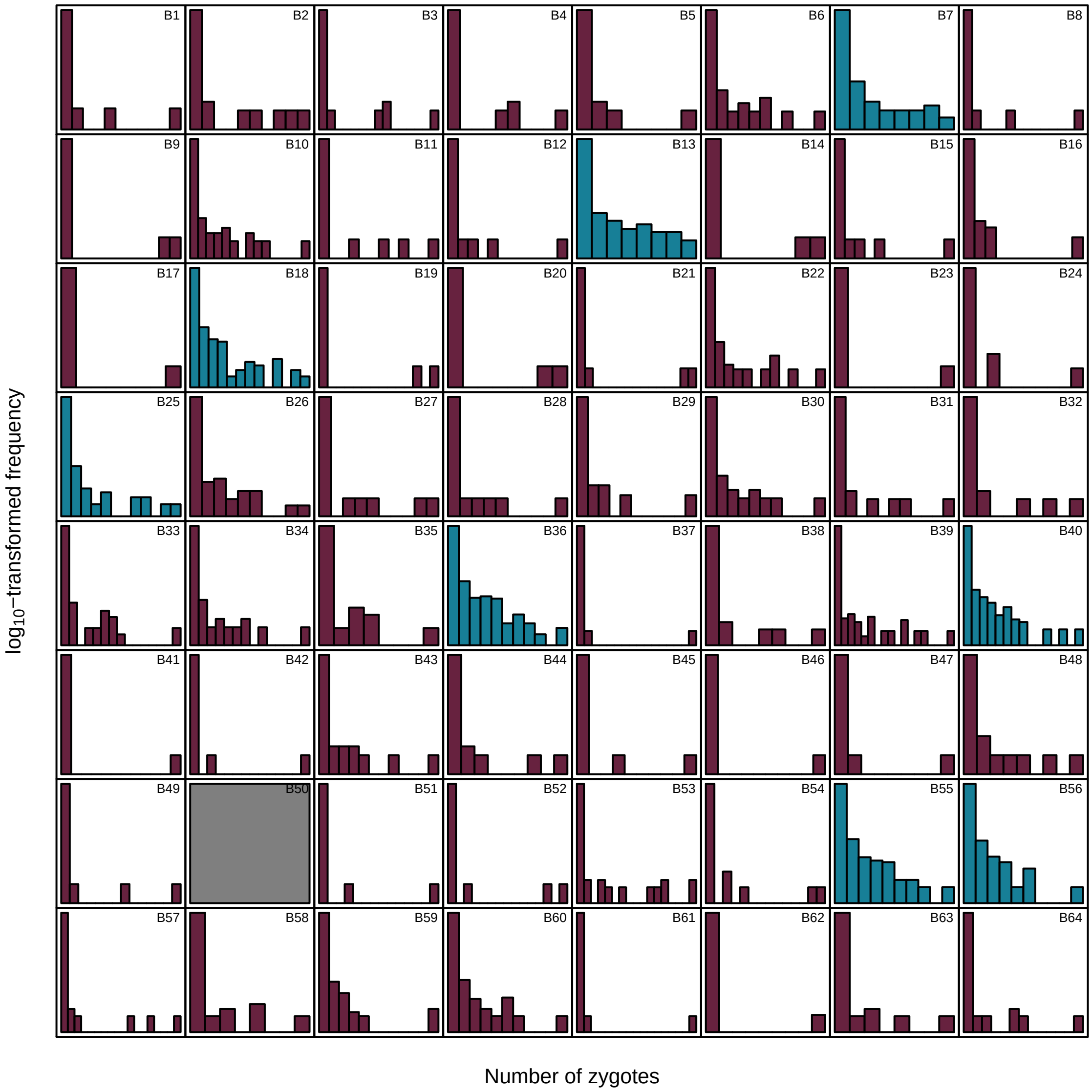


**Fig. S5.** Histograms of the number of zygotes produced per individual planktonic foraminifer across all experiments. Experiments resulting in low dispersion shown in teal, those with high dispersion shown in burgundy (compare article for details). Experiment B 50 showed no successful reproductive events.

### 2 Protist survival

In addition to the shell size data from planktonic Foraminifera presented in the manuscript, we used representative chamber-number counts from benthic Foraminifera to calculate comparable survival rates. We used data from PANGAEA (Lorenz 2005a, Lorenz 2005b, Foster et al. 2013a, Foster et al. 2013b, Foster et al. 2013c, Foster et al. 2013d, Foster et al. 2013e, Schmidt et al. 2018a, Schmidt et al. 2018b, Schmidt et al. 2018c), where chamber numbers in three different species of benthic Foraminifera had been counted systematically. We extrapolated the data to the initial population of one-chambered specimens (zygotes) and assumed that specimens with 15-chambers would be able to reproduce (Sen Gupta 2002). Using an exponential model, we could thus estimate survival rates in the benthic foraminiferal community (Fig. S6), which seems to be largely comparable with the survival of planktonic Foraminifera.

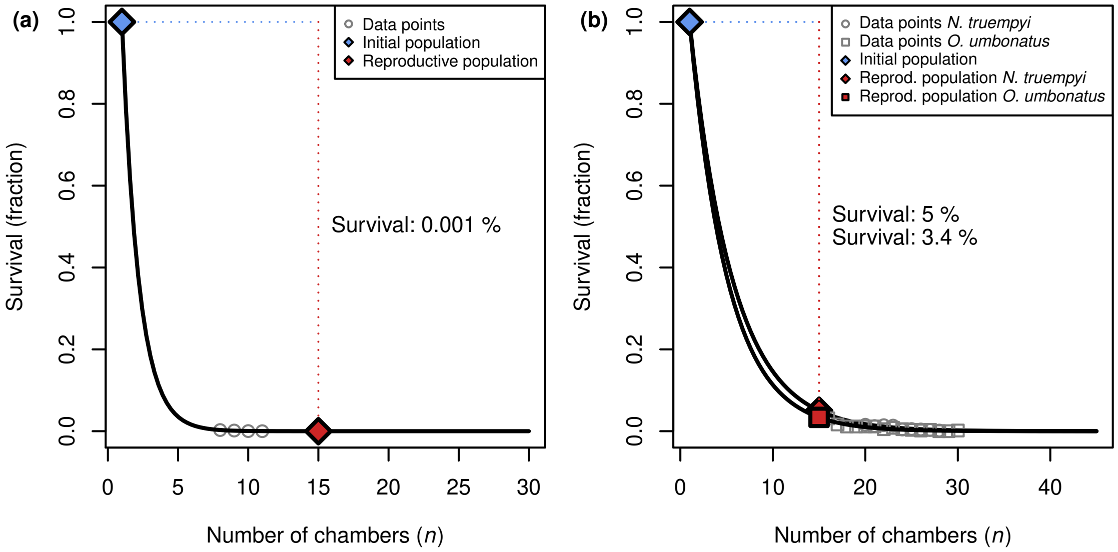


**Fig. S6.** Estimated survival curves for benthic Foraminifera based on representative chamber-number counts from the zygote (1 chamber) to the reproductive stage (15 chambers). (a) Data from *Cibicidoides wuellerstorfi* from the Denmark strait (Lorenz 2005a) imply very low survival rates, which may hint to some problems in the data. (b) Data from *Nuttallides truempyi* from the Palaeocene–Eocene (Foster et al. 2013c, Schmidt et al. 2018a) (survival rate: 5.0 %) and *Oridorsalis umbonatus* from the Palaeogene (Foster et al. 2013a, Foster et al. 2013b, Foster et al. 2013d, Schmidt et al. 2018b) (survival rate: 3.4 %) are very comparable to what we observed in planktonic Foraminifera.
